# DeepSNEM: Deep Signaling Network Embeddings for compound mechanism of action identification

**DOI:** 10.1101/2021.11.29.470365

**Authors:** C. Fotis, G. Alevizos, N. Meimetis, C. Koleri, T. Gkekas, LG. Alexopoulos

## Abstract

**Motivation:** The analysis and comparison of compounds’ transcriptomic signatures can help elucidate a compound’s Mechanism of Action (MoA) in a biological system. In order to take into account the complexity of the biological system, several computational methods have been developed that utilize prior knowledge of molecular interactions to create a signaling network representation that best explains the compound’s effect. However, due to their complex structure, large scale datasets of compound-induced signaling networks and methods specifically tailored to their analysis and comparison are very limited. Our goal is to develop graph deep learning models that are optimized to transform compound-induced signaling networks into high-dimensional representations and investigate their relationship with their respective MoAs.

**Results:** We created a new dataset of compound-induced signaling networks by applying the CARNIVAL network creation pipeline on the gene expression profiles of the CMap dataset. Furthermore, we developed a novel unsupervised graph deep learning pipeline, called deepSNEM, to encode the information in the compound-induced signaling networks in fixed-length high-dimensional representations. The core of deepSNEM is a graph transformer network, trained to maximize the mutual information between whole-graph and sub-graph representations that belong to similar perturbations. By clustering the deepSNEM embeddings, using the k-means algorithm, we were able to identify distinct clusters that are significantly enriched for mTOR, topoisomerase, HDAC and protein synthesis inhibitors respectively. Additionally, we developed a subgraph importance pipeline and identified important nodes and subgraphs that were found to be directly related to the most prevalent MoA of the assigned cluster. As a use case, deepSNEM was applied on compounds’ gene expression profiles from various experimental platforms (MicroArrays and RNA sequencing) and the results indicate that correct hypotheses can be generated regarding their MoA.

**Availability and Implementation:** The source code and pre-trained deepSNEM models are available at https://github.com/BioSysLab/deepSNEM.

**Contact:** Email for correspondence.

**Supplementary information:** Accompanying supplementary material are available online.

## 1 Introduction

Characterizing a compound’s Mechanism of Action (MoA) in a cellular system is a very important step in the development of new drugs or the repurposing of existing ones. On this front, several systems-based computational methods that utilize omics data, following treatment with a compound, have been developed [1]. One approach that has gained considerable attraction for the MoA identification task is the analysis of post-transcriptional data from compound perturbations [2]. These approaches analyze compounds’ transcriptomic signatures in order to identify key genes and signaling mechanisms that either cause the compound’s therapeutic effect or are associated with specific adverse effects [3]. Furthermore, the comparison of transcriptomic signatures can be used to elucidate the MoA of new compounds, by associating them with compounds of known MoA, or propose new indications for already existing drugs.

There have been many studies that utilize differential gene expression (GEx) data to characterize a compound’s MoA [4]. The Connectivity Map (CMap) and the LINCS project have played a pivotal role in this field, by providing large datasets of compounds’ transcriptomic signatures and methods for their analysis, comparison and interpretation [5,6]. As an example, Iorio et al. utilized compounds’ transcriptomic signatures from the CMap dataset to build a network, where perturbations are connected if they have similar transcriptional profiles [7]. This network was then analyzed to find communities and clusters that consisted of perturbations with similar MoA. Since a compound’s phenotypic effect is usually caused by changes in the expression of interacting genes/proteins, combining transcriptomic data with a prior knowledge-base of molecular interactions, e.g. signaling pathways, can result in a more mechanistic explanation of a compound’s MoA [1]. On this front, a promising modeling technique is the representation of a compound’s effect as a network of signaling proteins (nodes), showing their activity and how these interact with each other to transfer the signal of the perturbation in the system [8].

Signaling network creation methods combine omics data with a prior knowledge network of protein-protein interactions (PPI) in order to extract a graph that best explains the experimental data. Mitsos et al. developed an Integer Linear Programing (ILP) optimization task to identify the signaling network that characterizes a compound’s effect based on phosphoproteomic data [9]. Since large scale phosphoproteomic datasets following compound treatment are very rare, there has been a concentrated effort to develop methods for signaling network creation based on transcriptomics [10-12]. Liu et al. developed CARNIVAL, a causal reasoning framework to identify signaling networks that best explain a set of transcription factor (TF) activity scores, calculated from differential GEx data [13]. Compound-induced signaling networks are information-rich and complex representations of the compounds’ effect, since they incorporate the prior knowledge of molecular interactions in the form of a PPI network. However, this complexity poses limitations for their large scale analysis and comparison of networks from different compounds using traditional network similarity algorithms, i.e. graph kernels. More specifically, graph similarity algorithms, such as the Graph Edit Distance (GED), graphlet-based methods or graph kernels, utilize hand crafted features and are not optimized for signaling networks, which can result in reduced generalization performance and reduced scalability [14-16]. An interesting approach is to employ deep learning models for graphs in order to encode the complete information of the signaling network into high dimensional fixed-length representations [17]. These representations can then be compared using traditional algorithms in order to identify similarities between compound-induced signaling networks that could translate to similarities in the compounds’ MoA.

There have been many studies for the development of deep learning models for graph data in a variety of fields. These models are usually neural networks that aim to learn new task-specific node and graph representations by using the graph’s connectivity [18]. For example, the graph convolutional model utilizes a message passing algorithm to learn neighborhood-level representations of the input graph. Recently, the successful transformer architecture for natural language processing (NLP) problems has been modified and applied on graph data [19,20]. Graph transformers utilize an attention mechanism for each node that is a function of the neighborhood’s connectivity, rather than a message passing algorithm. Similarly, the graph2vec model was inspired by the doc2vec approach for NLP tasks. Graph2vec treats the entire graph as a document and each node’s neighborhood as a word and aims to learn a fixed-length representation of the entire graph in a fully unsupervised task [21]. Another important unsupervised approach for graph representation learning is the InfoGraph model [22,23]. InfoGraph aims to maximize the mutual information between graph-level representations and representations of the graph’s substructures at different levels, e.g. nodes, edges and triangles. These unsupervised graph representation learning methods can be modified for compound-induced signaling networks in order to extract fixed-length feature vectors that can then be associated with the compound’s MoA.

In this paper, we developed a novel deep learning framework, called deepSNEM, to learn new representations (embeddings) of signaling networks and investigate their relationship with the compound’s MoA. Compounds’ signaling networks were created using the CARNIVAL pipeline and the transcriptomic signatures of the CMap dataset, resulting in a large scale dataset of signaling networks that can aid future studies. The core of deepSNEM is an unsupervised graph transformer trained to maximize the mutual information between representations of graphs’ substructures that belong to signaling networks created from similar perturbations. The resulting embeddings were evaluated based on their ability to identify similar signaling networks and compared with representations created by different graph-based models. Subsequently, the embeddings were clustered with the k-means algorithm and the resulting clusters were analyzed based on their MoA composition. Furthermore, a subgraph importance method was developed to identify the most important nodes for each graph-level representation and the subgraphs that cause the signaling networks to cluster together. As a use case, deepSNEM was tasked to assign clusters to compounds’ signaling networks generated using gene expression profiles from various experimental platforms. Analyzing the MoA composition of a compound’s assigned cluster, deepSNEM can generate hypotheses regarding the MoA of new lead compounds or suggest new potential mechanisms for already existing drugs.

## 2 Results

### 2.1 The deepSNEM approach

The overview of our approach is presented in Figure 1. Differential gene expression signatures following compound treatment across cell lines were retrieved from the L1000 dataset (GSE92742) [6]. In total, 7722 signatures from 3005 compounds across 70 cell lines were utilized. The first step of the deepSNEM pipeline is the creation of signature specific signaling networks following the CARNIVAL framework [13]. In this framework, the gene expression signatures are first transformed into transcription factor activity scores and then an ILP model is tasked to extract the optimal subgraph from a global PPI network that best fits the calculated activity scores (see Methods 4.1). The created network is a labeled (protein activity), signed (edge activation or inhibition) and directed PPI graph that captures the signaling network effect of the drug-induced transcriptomic signature. The core of deepSNEM is a DL model, trained in an unsupervised setting, which takes as input the drug-induced signaling networks, created with CARNIVAL, and outputs a high dimensional embedding that best captures the information contained in the input graph. Regarding the DL models, we evaluated the use of a graph transformer trained to either maximize the mutual information of nodes belonging to the same signature (termed deepSNEM-GT-MI) or predict the edge presence between nodes (termed deepSNEM-GT-LP), a siamese GCN model to predict the graph edit distance between signaling networks (termed deepSNEM-GED) and the widely used graph2vec model (termed deepSNEM-G2V) (see Methods 4.2).

**Figure 1.**
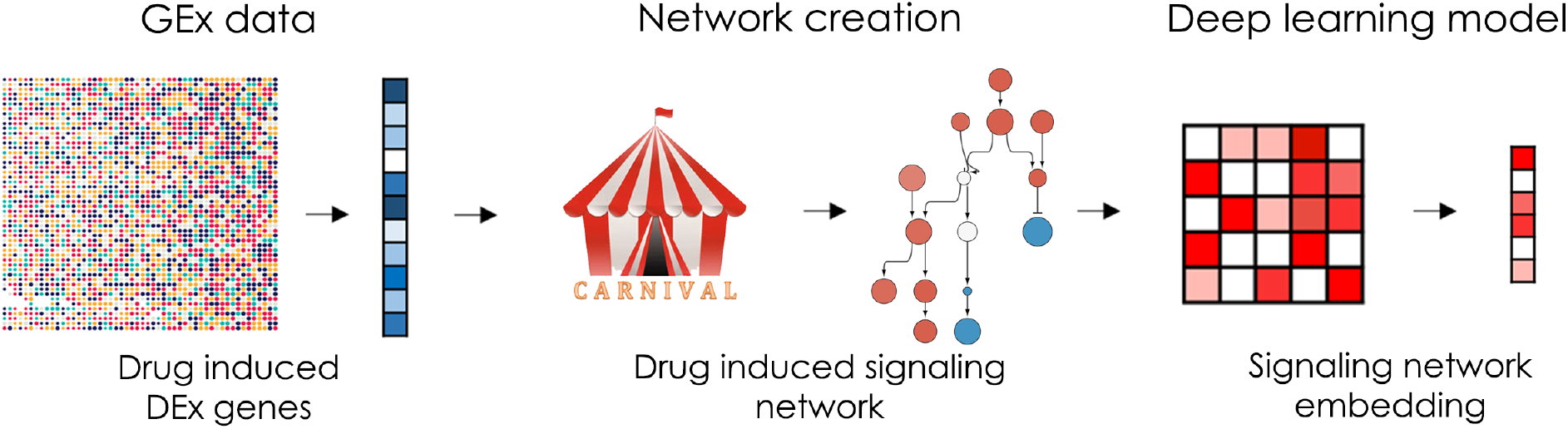
Schematic overview of deepSNEM. For each compound-induced differential expression signature, a signaling network is created using the CARNIVAL framework. Then an unsupervised DL model is tasked to encode the created signaling network in a high dimensional embedding that best captures the input graph information.

### 2.2 Model-embedding evaluation

The different deepSNEM model variations were evaluated based on the validity of the produced embeddings on two separate tasks. The first task examines the models’ ability to produce similar embeddings from signaling networks that are created from the same differential gene expression signature. On this front, we utilized the slightly different but feasible network solutions of CARNIVAL’s ILP model for the same signature and investigated the distributions of Euclidian distances between embeddings belonging to the same signature and between embeddings from different signatures (Figure 2A). As it can be seen in Figure 2A, there is a clear distinction between the distance distributions of embeddings from the same and different signatures. Thus, all models are able to produce embeddings that are significantly more similar for graphs created from the same measurements of differential expression. In the second task, we evaluated the similarity of graph embeddings created from duplicate gene signatures as compared to the similarity of embeddings from random gene signatures. Duplicate signatures indicate transcriptomic signatures from the same compound perturbation, cell line, dose and time point that were assayed on different L1000 plates [24]. Figure 2B shows the distributions of Euclidian distances between embeddings belonging to duplicate signatures and between embeddings of random signatures. For all models, the difference between the distributions is significant, as indicated by a two sample t-test (p-values < 0.001). Thus, all models are able to produce similar graph embeddings for gene signatures that share the same experimental conditions. Based on these results, we chose to perform a clustering analysis on the embeddings produced by the deepSNEM-GT-MI architecture, in order to examine the connection between a drug’s induced signaling network and its reported MoA.

**Figure 2.**
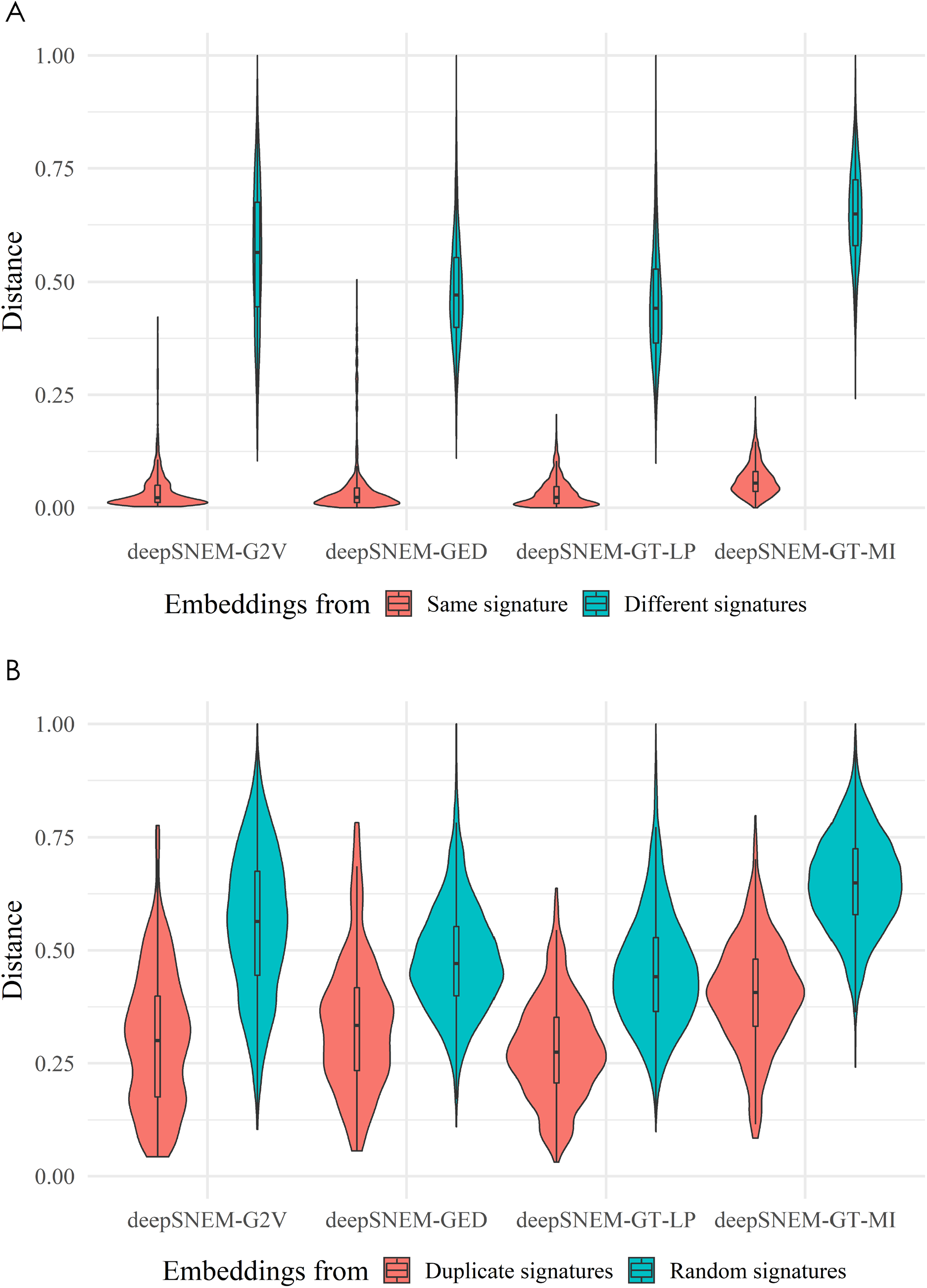
Model-embedding evaluation tasks. In the deepSNEM-G2V model the graph2vec architecture is used to embed the graphs into the latent space. The deepSNEM-GED approach is a distance learning approach, where the model is trained so that the Euclidian distance of the embeddings corresponds the graph edit distance of the
signaling networks. The deepSNEM-GT-LP architecture is a graph transformer-based encoder trained on the edge prediction task. Finally, the deepSNEM-GT-MI is the graph transformer-based encoder trained to maximize the mutual information between the embeddings of similar graphs. (A) Normalized Euclidian distances between embeddings from the same signature and different signatures for all deepSNEM model variations. (B) Normalized Euclidian distances between embeddings duplicate and random gene expression signatures for all model variations.

### 2.3 Clustering analysis for MoA identification

The signaling network effect of a compound perturbation in a cellular model presents a systematic view into the compound’s MoA. In order to investigate this relationship, we first identified groups of perturbations with similar network effect, by clustering the deepSNEM network embeddings, and then analyzed the resulting clusters based on the reported MoA of the compounds. On this front, the 256-dimensional deepSNEM-GT-MI embeddings were clustered using the k-means algorithm. The optimal number of clusters was found to be 200, according to the k-means elbow plot (see Supplementary Material (SM) 6). Additionally, in order to analyze and characterize the resulting clusters, we utilized the MoA labels provided by the Broad’s Institute Repurposing Hub [25]. Out of the 3005 unique compounds, 912 were mapped to 261 unique MoA labels using the Repurposing Hub dataset (see SM 1). Figure 3A shows the 2-dimensional t-SNE projections of all available signaling network embeddings. Additionally, the signaling network embeddings that belong to the top 9 most prevalent MoA labels in the dataset are presented with different colors (Figure 3A). In order to characterize the identified clusters, we focused on the subset of clusters that are significantly enriched for at least one mechanism (Figure 3B). The selected clusters have at least 25% of their compound perturbations belonging to the same MoA, with a p-value lower than 10^−6^ compared to a random selection. Figure 3B shows the breakdown of the available MoA in the selected clusters. As it can be seen, the identified clusters are enriched for the same mechanisms that are most prevalent in the labeled dataset. As a result, DeepSNEM was able to identify 11 clusters that are significantly enriched for specific mechanisms, i.e. mTOR, HDAC, topoisomerase, protein and ATP synthesis inhibitors. We have to note that clusters that are enriched for MTOR inhibitors are also enriched for PI3K inhibitors, which is expected due to the PI3K/mTOR signaling pathway. However, the majority of the compounds in each cluster still do not have available labels regarding their MoA (represented with grey color in Figure 3B). Thus, due to the unknown labels, the distribution of MoA between clusters that are enriched for the same MoA can still be quite different.

**Figure 3.**
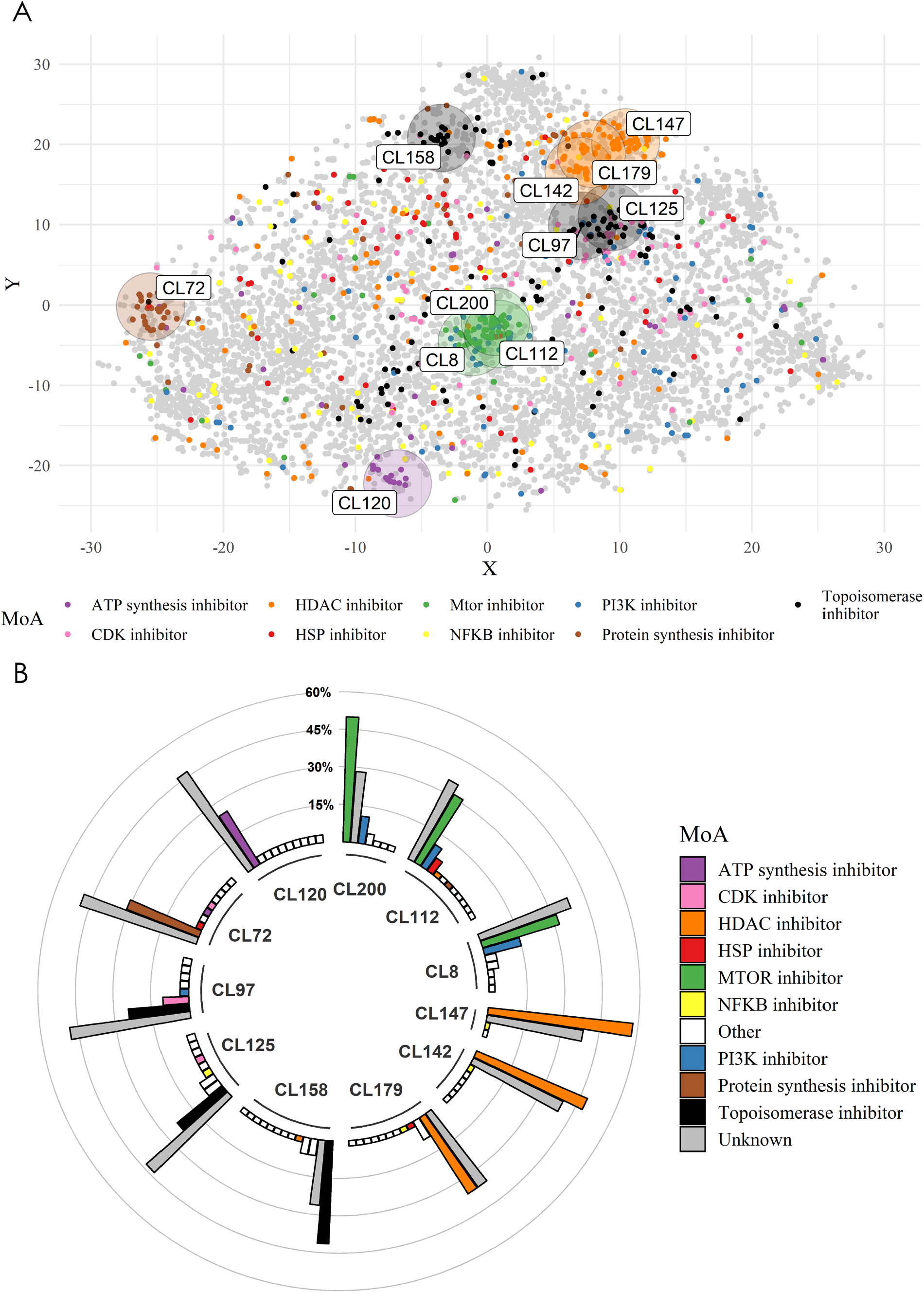
Clustering analysis. (A) T-SNE projection of the 256-dimensional signaling network embeddings of deepSNEM-GT-MI. Different colors represent the 9 most prevalent MoA in the dataset, while the grey color represents perturbations with either unknown or other MoA. Additionally, the centers of the identified clusters are represented with circles (CL: cluster). (B) MoA composition of the analyzed clusters. The Y axis represents the frequency, as a percentage, of each MoA in the cluster (CL: cluster).

### 2.4 Subgraph importance

The analysis of compound-induced signaling networks for MoA identification offers the benefit of easier result interpretation. In order to utilize this benefit and increase the interpretability and explainability of deepSNEM, we created a framework to identify the important subgraphs for the subset of clusters analyzed in the previous section. For each cluster, important nodes were identified using an aggregate score based on their importance to the embedding model and the nodes’ prevalence in the cluster’s graphs (see Methods 4.3). Figure 4A shows the overlap, as a percentage, between the 20 most important nodes of the analyzed clusters. As it can be seen, clusters that are enriched for the same MoA, have a higher similarity between their most important nodes. Thus, the proposed importance framework can identify nodes of high importance in each cluster that show a connection to the cluster’s most prevalent mechanism of action. For visualization purposes, the most important nodes in each cluster were connected by selecting the shortest paths between them, from the Omnipath PPI that also maximize the overall sum of importance scores in the path. Figure 4B shows an example of the important subgraphs for the clusters that are enriched for mTOR and PI3K inhibitors. The common most important nodes across the presented networks include the mTOR regulated transcription factors NRF1 and TFDP1 and the CSKNK2A1, RHOA, PRKACA and LCK proteins, which are involved in the PI3K-Akt-mTOR signaling pathway [26-30]. Finally, across all clusters, AKT1 and MAPK1 serve as central nodes that connect the most important nodes (Figure 4B). The important subgraphs for all analyzed clusters are presented in SM 7.

**Figure 4.**
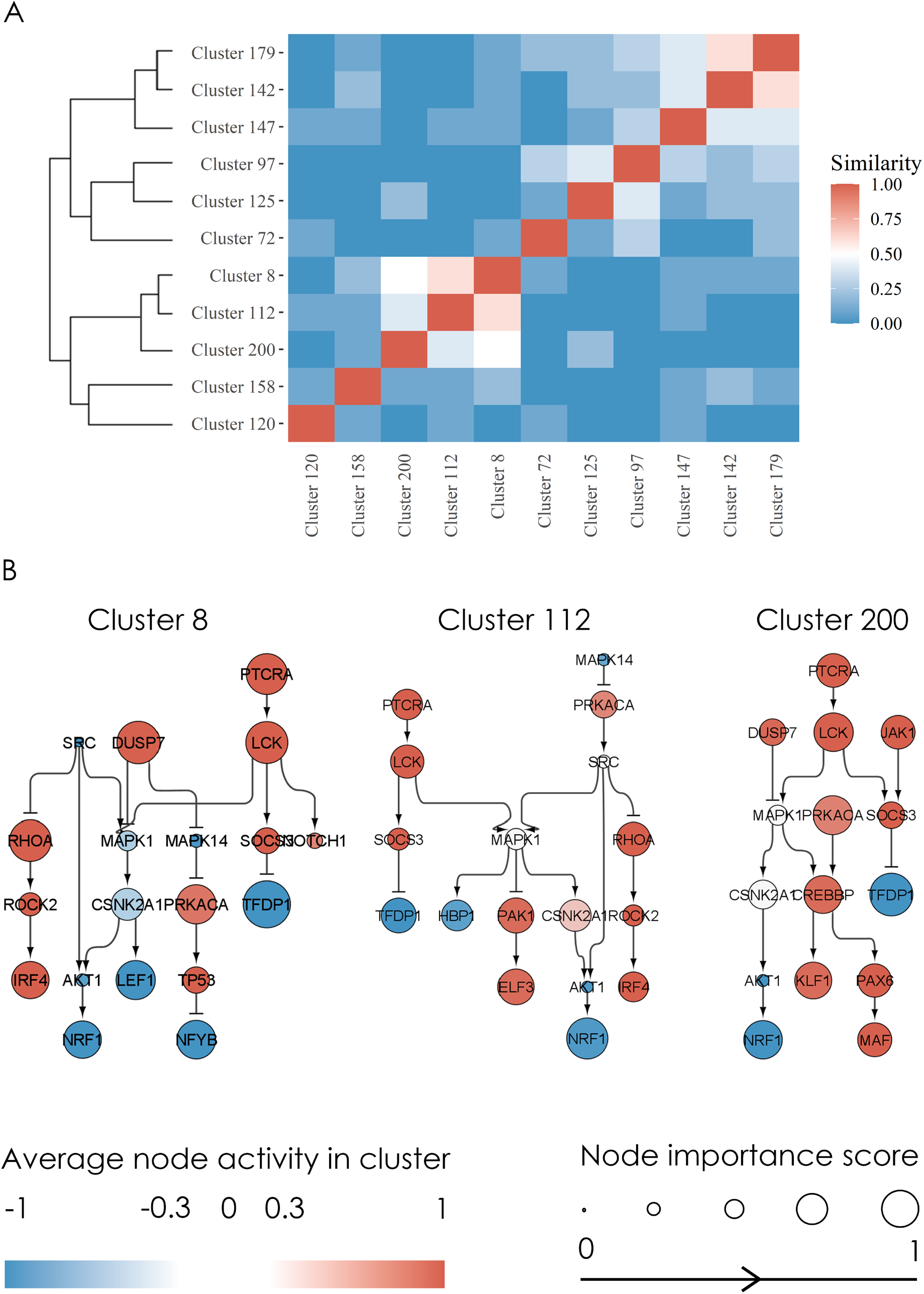
Cluster subgraph importance. (A) Heatmap showing the similarity, as percentage overlap, between the 20 most important nodes of each cluster. (B) Important subgraphs identified for the clusters enriched for mTOR and PI3K inhibitors (Clusters 8, 112 and 200). The average activity of each node in the cluster is color coded from blue to red. Blue nodes are inhibited, while red are activated. Each node’s importance score, ranging from 0 to 1, is represented by the size of the node’s circle.

### 2.5 Use case: cluster assignment

Gene expression data from 7 additional compounds with known mechanism of action were retrieved from the GEO database. The details regarding the experimental data used in the use case are presented in Table 1. Overall, the data were collected from 6 different studies, 4 cell lines and 3 different experimental platforms, i.e. Affymetrix/Agilent Microarrays and Illumina next generation sequencing. Following the deepSNEM pipeline, each differential gene expression signature was transformed into a compound induced signaling network with CARNIVAL and embedded using the deepSNEM-GT-MI model. Finally, each embedding was assigned to one of the already identified clusters (Table 1). Figure 5A shows the assigned clusters and the distribution of each cluster’s available MoA. The topoisomerase inhibitor SN38 and the HDAC inhibitors Sodium-Butyrate, Panobinostat and Belinostat were assigned to clusters significantly enriched for topoisomerase and HDAC inhibitors respectively. Furthermore, the topoisomerase inhibitor Doxorubicin and the mTOR inhibitor Sirolimus were assigned to clusters enriched for their respective MoA, albeit having a large number of compounds with unknown MoA. Finally, the compound CDK-887 was assigned to a cluster that was not enriched for any particular MoA. Thus, the deepSNEM pipeline can be used to assign a cluster to a compound-induced gene expression signature, independent of the experimental platform, and provide insight into the compound’s potential MoA. For the compounds in the use case, we also compared the cluster assignment of deepSNEM to a clustering of the compounds’ differential expression gene measurements into the same number of clusters (k=200) (see SM 8) (Figure 5B). Comparing the two approaches, SN38, Belinostat and Panobinostat were assigned to clusters composed of similar mechanisms. However, this is not the case for Sirolimus, Doxorubicin and Sodium Butyrate, which are assigned to clusters not enriched for any particular MoA, when the gene-clustering pipeline is used. Finally, for each compound of the use case, we calculated the Jaccard similarity index between the perturbations of the identified clusters using the two methods (deepSNEM and gene-based clustering) (Table 2). As it can be seen in Table 2, across all compounds the similarity of the clusters is very low, with only the clusters assigned to the SN38 having a slightly higher Jaccard index. Thus, the deepSNEM and gene-based pipeline result in a different clustering of the perturbations, due to the different biological hierarchy of information provided by the compound-induced signaling networks and differential gene expression signatures.

**Table 1.** Information regarding the perturbations used in the use case and their assigned clusters.

| Compound | MoA | Cell line | GSE | Platform | Cluster (CL) |
| --- | --- | --- | --- | --- | --- |
| Sirolimus | mTOR inhibitor | MCF7 | GSE116447 | Affymetrix Microarray | 53 |
| CDK-887 | CDK inhibitor | MCF7 | GSE19638 | Affymetrix Microarray | 163 |
| Panobinostat | HDAC inhibitor | A375 | GSE145447 | Illumina<br>NextSeq | 22 |
| Sodium-Butyrate | HDAC inhibitor | HT29 | GSE61429 | Agilent<br>Microarray | 22 |
| Belinostat | HDAC inhibitor | A549 | GSE96649 | Illumina<br>NextSeq | 188 |
| SN38 | Topoisomeras<br>e I inhibitor | MCF7 | GSE18552 | Affymetrix<br>Microarray | 158 |
| Doxorubicin | Topoisomeras<br>e II inhibitor | MCF7 | GSE19638 | Affymetrix<br>Microarray | 33 |

**Table 2.**
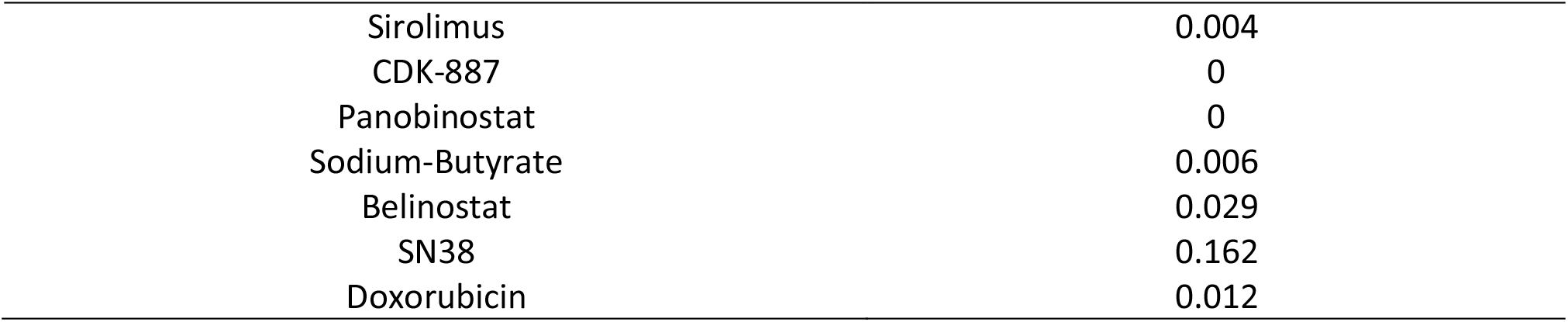
Jaccard similarity index between the clusters that the use case compounds were assigned to, using the gene-based and deepSNEM pipelines.

|  |  |
| --- | --- |
| Sirolimus | 0.004 |
| CDK-887 | 0 |
| Panobinostat | 0 |
| Sodium-Butyrate | 0.006 |
| Belinostat | 0.029 |
| SN38 | 0.162 |
| Doxorubicin | 0.012 |

**Figure 5.**
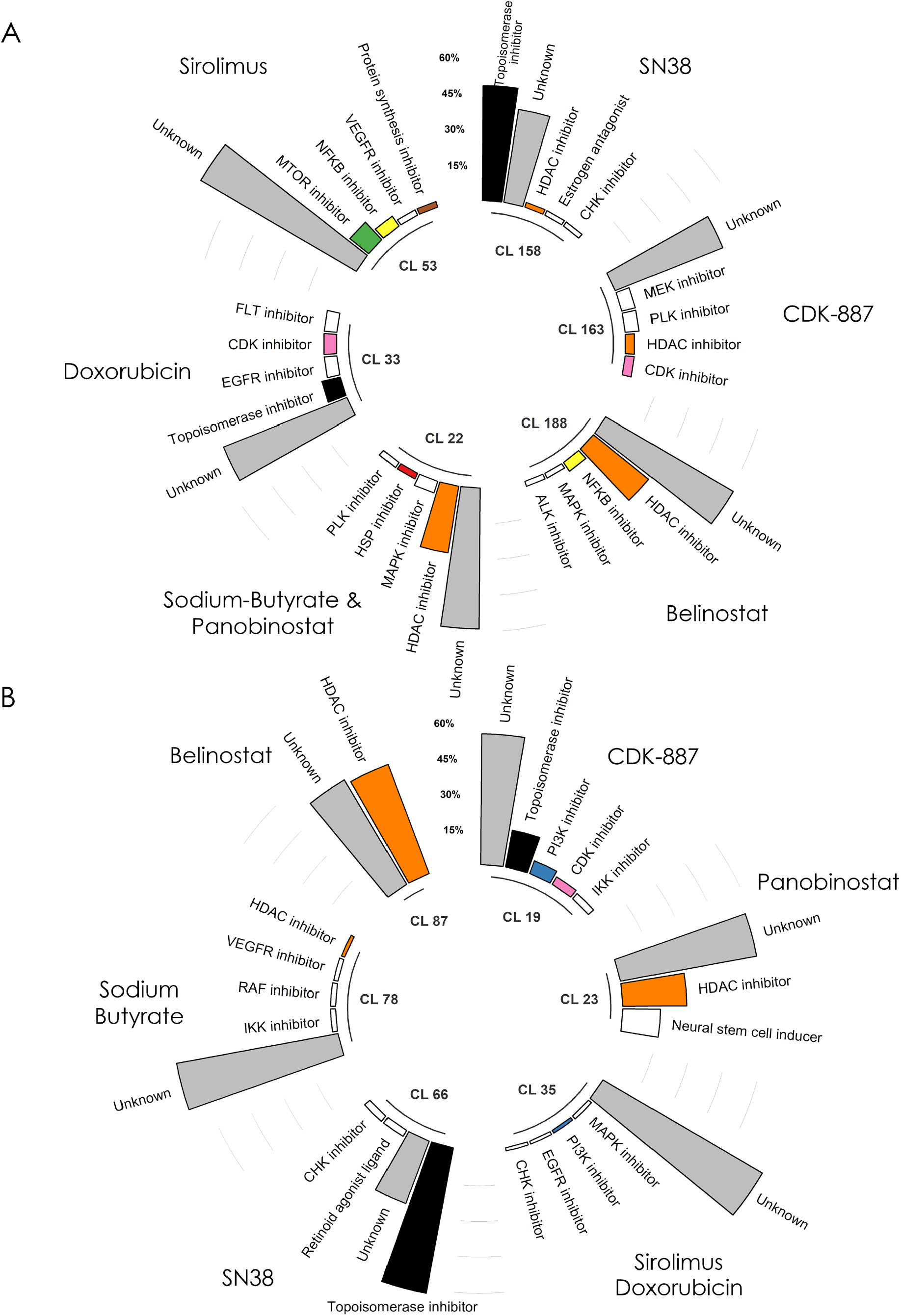
MoA composition of the compounds’ clusters. (A) Bar plot of mechanism of action prevalence for the clusters that were assigned to the use case perturbations using the deepSNEM pipeline. (B) Similar bar plot for the assigned clusters using the gene-based clustering pipeline.

## 3 Discussion

The changes in the protein signaling network caused by a compound perturbation can aid in studying the compound’s mechanism of action in the cellular system. However, analyzing compound-induced signaling networks on a massive scale is a very complex problem, not only due to the limited availability of large datasets containing such networks but also due to the complex structure of the data. This complex structure of signaling networks limits their representation abilities and poses a challenge in identifying similarities or differences between them. In this study, we created a large dataset of compound-induced signaling networks from the CMap dataset, using the CARNIVAL network creation pipeline and developed an unsupervised deep learning model to transform them into high-dimensional and information-rich representations. This novel approach, called deepSNEM was used to identify clusters of perturbations with similar network representations and offer insight into the compounds’ MoA by analyzing the distribution of MoA in the clusters.

The prediction of a compound’s MoA from biological response data has gained considerable attraction in the machine learning community [31,32]. This is evident by the recent release of the CTD^2^ Pancancer Drug Activity DREAM Challenge, which tasked the community to predict a compound’s MoA based on post-transcriptional and cell viability data [32]. Even though the learning task of MoA prediction is frequently modeled as supervised, in our approach we decided to develop deepSNEM in a fully unsupervised fashion. This decision was based on the nature of the learning task and the compounds’ MoA, wherein if a compound has a reported MoA based on binding affinity data, we can’t know with absolute certainty that it doesn’t have additional MoA labels due to other binding targets or interactions between the proteins in a pathway. Thus, for some compounds the negative labels for all possible MoA indications might not be truly negative, rather they might be simply unknown. Additionally, another important benefit of using an unsupervised approach, is that we can greatly increase the amount of available data by including transcriptomic signatures following treatment with compounds that have no reported MoA. In deepSNEM the learning model is tasked to produce meaningful representations that capture the information included solely in the compound-induced signaling networks without taking into account the compounds’ reported MoA. However, this unsupervised task makes the evaluation of the different models and the resulting embeddings quite challenging.

The evaluation of the validity of the resulting embeddings was based upon two tasks that test if the models can produce embeddings that capture the similarities of the input perturbation. Those tasks however, more closely resembling pass/fail tasks, rather than quantitative metrics (Figure 2). Thus, we cannot know with certainty which deepSNEM model variation, i.e. graph transformers, graph convolutions or graph2vec is better in terms of the resulting embeddings. For the downstream task of mechanism of action identification, we decided to use the embeddings of the graph transformed trained to maximize the mutual information between nodes that belong to networks created from the same or duplicate gene expression signatures. We argue that this deepSNEM variation is better suited to capture the information of the signaling networks, due to the graph transformer architecture and due to the mutual information task that forces networks created from the same perturbation to have similar embeddings (see Methods 4.2). Finally, we have to note that the resulting 256-dimensional graph embeddings contain all the information of the input signaling networks, which makes it difficult for the t-SNE algorithm to project them in 2 dimensions, as it can be seen in Figure 3A.

The clustering analysis and MoA identification using the deepSNEM-GT-MI embeddings was performed by analyzing the MoA labels provided by the Broad Institute in the drug repurposing hub. Using this dataset, 912 out of the 3005 total compounds were mapped to 261 unique labels. We argue that this diversity of mechanisms and large number of compounds with unknown MoA in the dataset resulted in the large number k (k = 200) of clusters that were identified using the elbow plot of the k-means algorithm. Additionally, due to the large number of unlabeled compounds, in order to analyze the resulting clusters, we focused on a specific subset that is significantly enriched for at least one specific MoA (Figure 3B). Using this approach, we identified 11 clusters that each were enriched for the most prevalent mechanisms in the dataset. However, even for the clusters enriched for the same MoA, the large number of unknown compounds could result in different cluster compositions, which potentially further signifies the importance of analyzing biological response from different points of view, e.g. genes, pathways, signaling networks.

There have been many studies for the identification of a compound’s MoA using biological response data. The majority of these approaches utilize post-transcriptional data and have been utilized successfully in the fields of systems pharmacology and drug repurposing [34,35]. Since the initial part of deepSNEM relies on transcriptomic data, similarities between the results and clustering of gene signatures and signaling networks are expected. This effect is evident in the presented use case, where some of the compounds were assigned to clusters with similar MoA composition between the gene-based and network-based pipeline. However, some compounds were assigned to clusters enriched for different MoA between the two approaches (Figure 5). Most importantly, between the two methods, each compound was assigned to clusters that had a very low Jaccard similarity index, meaning that the transcriptomic signatures and signaling network embeddings of deepSNEM cluster in a different way (Table 2). Thus, even though transcriptomic signatures do provide meaningful insight into a compound’s MoA, there are cases, where analyzing the signaling networks can reveal complex relationships that are hidden in the original expression data. We argue that this is because a compound’s effect on a biological system is usually caused by changes in the expression of genes that interact with each other to form specific biological processes. By supplying deepSNEM with this required prior knowledge of interactions in the form of the Omnipath PPI, the compound-specific signaling networks can provide a mechanistic view of the compound’s effect and translate to the identification of its MoA [36]. Additionally, deepSNEM’s signaling network creation via the CARNIVAL pipeline can provide a robust normalization factor to analyze and incorporate data from different experimental platforms (Table 1). Finally, the analysis of compound-induced signaling networks has the inherent benefit of increasing the interpretability of results.

The interpretability and explainability of machine learning models is a concept that has gained considerable attraction since the creation and application of powerful and complex deep learning models in various fields [37]. This is especially true in the fields of drug discovery and systems pharmacology, where understanding why the model made specific decisions and predictions can not only validate and help interpret the results, but also generate new knowledge and hypotheses regarding the complex systems under study [38]. Here, we developed a node and subgraph importance method to identify which nodes the model pays attention to when creating the embeddings and which nodes in the original networks cause the embeddings to cluster together. This resulted in the better understanding and interpretation of the novel representations that were extracted from the DL model. Using this approach, we showed that the models pay attention to similar nodes in order to cluster together compounds with similar MoA and were able to identify important signaling subgraphs that are characteristic of each cluster (Figure 4). For example, in the clusters enriched for mTOR inhibitors, even though mTOR as a node was not present in the input signaling networks of the cluster, deepSNEM was able to extract important subgraphs that are related to the mTOR signaling pathway.

The deepSNEM pipeline serves as proof of concept that compound-induced signaling networks can be analyzed on a massive scale, using deep learning and provide insight into the compound’s effect. In a real-world application, deepSNEM would be used in combination with existing methods, utilizing transcriptomic data or pathway signatures, for a consensus-based assignment of compound perturbations into clusters that are enriched for specific MoA. Subsequently, deepSNEM could be used to identify which nodes and subgraphs mostly influenced the proposed cluster assignment, thus increasing its interpretability and help generate new hypotheses. Finally, deepSNEM could be combined with different signaling network creation. We believe that our signaling network dataset and the proposed pipeline can help pave the way towards more studies that utilize the inherent knowledge of the changes in the signaling cascade of a system to better elucidate a compound’s mechanism of action.

## 4 Methods

### 4.1 Signaling network creation

Gene expression profiles (level-5 z-score transformed) of compound perturbations were downloaded from the L1000 CMap dataset [6]. In the current study, only measurements of the relative gene expression of the 978 landmark genes in the L1000 assay were used (GSE92742). For each gene expression signature, a quality score was derived, based on its transcriptional activity score (TAS), the number of biological replicates and whether the signature is considered an exemplar, similar to the deepSIBA approach [24]. Based on this quality score, only the signatures with the highest quality score were selected. An overview of the transcriptomic signatures used in this study can be found in SM 1. For each signature, transcription factor (TF) activity scores were inferred using the DoRothEA R package [39]. This method utilizes a knowledge base of signed TF-target interactions called Regulons and the VIPER enrichment algorithm to calculate TF activity scores [40]. For each compound perturbation, the discretized TF activities of DoRothEA were transformed into signaling networks using the CARNIVAL pipeline [13]. CARNIVAL solves an ILP optimization problem to infer a family of highest scoring subgraphs, from a prior knowledge network of signed and directed protein-protein interactions, which best explain the TF activities, subject to specific constraints. In our approach the OmniPath network was used as the global prior knowledge network [36]. Furthermore, the CARNIVAL pipeline without using the perturbation targets as input was utilized (InvCARNIVAL method). Finally, the ILP formulation of the problem was solved using the IBM ILOG CPLEX solver, which is freely available through the Academic Initiative (https://www.ibm.com/products/ilog-cplex-optimization-studio). Details regarding the parameters of CARNIVAL can found in SM 2.

### 4.2 DeepSNEM model

#### 4.2.1 DeepSNEM-GT-MI

Each compound-induced signaling network is represented as a labeled, signed and directed graph *G* = (*V,E*), with nodes (V) being the proteins and edges (E) denoting the directed physical interaction between them. Additionally, the activity of each protein is represented as a node attribute, while the inhibition or activation of each edge is represented as an edge attribute. Each input graph to the deepSNEM-GT-MI consists of a node feature matrix (X_prot_) and a node activity embedding (X_act_). The node feature matrix contains the initial protein features of each graph, which were created using the SeqVeq protein sequence model [41]. For each protein, the node activity embedding is a projection of the node’s activity to the dimensions of the SeqVeq features, using a single embedding layer. The node feature and node activity matrices are added before being processed by the graph transformer. The input matrices are then passed through the self-attention mechanism of the graph transformer, resulting in a final feature matrix X [19,20]. Finally, this feature matrix is summarized using the Set2Set global pooling method into a trainable whole-graph representation [42]. The model is trained fully unsupervised by maximizing the mutual information between node and whole-graph embeddings that are created from the same or duplicate transcriptomic signatures, using the CARNIVAL pipeline, thus resulting in similar graph representations for the same perturbation. Similar to the InfoGraph approach, the Jensen-Shannon Mutual Information estimator was used, while an additional term was added to the total loss function in order to force the embeddings to be uniformly distributed [22]. More details regarding the deepSNEM-GT-MI model can be found in SM 5.

#### 4.2.2 DeepSNEM model variations

The DeepSNEM-GED variation is a Siamese graph convolutional model that is trained to minimize the error between the predicted and calculated graph edit distance for a pair of compound-induced signaling networks. Furthermore, the deepSNEM-GT-LP variation is a transformer model similar to deepSNEM-GT-MI, albeit trained to predict the presence of an edge between two proteins (nodes). Finally, the deepSNEM-G2V model is an application of the widely used graph2vec model for whole-graph representations [21]. Details regarding these model variations can be found in SM 3 and 4.

### 4.3 Node and subgraph importance

The average attribution of each node (protein) to the resulting signaling network embedding was calculated using the saliency map approach of the Captum library [43]. With the saliency approach the attributions are calculated based on the gradient with respect to the input [44]. This approach results in an attribution score for each node that shows the importance of the node to the model, when calculating the network embedding. Subsequently, a scoring function was designed in order to identify the important nodes in a specific cluster of signaling network embeddings. For each node, this scoring function calculates the product of the median rank of the node’s attribution score in the cluster and the frequency that the node appears in the signaling networks of the cluster. Finally, this score is normalized between 0 and 1. For visualization purposes, the 20 most important nodes of each cluster were connected using the shortest paths from the OmniPath PPI network that maximize the overall sum of importance scores in the connected graph.

## 5. Acknowledgements

We would like to thank Dr. Avlant Nilsson for their constructive feedback on the manuscript.

