## Supplementary Material for "DeepSNEM: Deep Signaling Network Embeddings for compound mechanism of action identification"

### 1 Data preprocessing and quality control

The filtered CMap dataset contains 7722 transcriptomic signatures from 3005 compounds tested across 70 cell lines. During the filtering process, for each compound per cell line, its signature with the highest quality across different dosages and time points was selected. The assigned quality score based on TAS, number of replicates and whether the signature is considered an exemplar is presented in Table S1. Only signatures with Quality score of 1 were used.

**Table S1** Signature quality score (re-created from Fotis et al. [5])

| Quality score | TAS | Number of replicates | Exemplar |
|---|---|---|---|
| Q1 | > 0.4 | > 2 | True |
| Q2 | 0.2 – 0.4 | > 2 | True |
| Q3 | 0.2 – 0.4 | ≤ 2 | True |
| Q4 | 0.1 – 0.2 | > 2 | True |
| Q5 | 0.1 – 0.2 | ≤ 2 | True |
| Q6 | < 0.1 | > 2 | True |
| Q7 | < 0.1 | ≤ 2 | True |
| Q8 | < 0.1 | < 2 | False |

### 2 CARNIVAL parameters

The CARNIVAL pipeline was ran in parallel and without using the perturbation's known targets as input (InvCARNIVAL). The signaling network dataset was created with an older version of CARNIVAL in R version 3.6, but the same parameters can be used in the latest version of CARNIVAL.

The main parameters, which can be found in Table S2, are the time limit until the optimization terminates (timelimit), the allowed number of solutions to be generated (limitPop), the allowed number of solution to be kept in the pool of solution (poolCap) and the external ILP Solver used. The rest parameters can be set to the default of each CARNIVAL version [1].

**Table S2** CARNIVAL pipeline parameters

| **Execution mode** | parallel |
|---|---|
| **inverseCR** | TRUE |
| **ILP Solver** | Cplex |
| **timelimit (in minutes)** | 1800 |
| **limitPop** | 500 |
| **poolCap** | 100 |

### 3 Graph2vec

Our approach was compared with a well-known and well-established model for the generation of graph embeddings, called graph2vec [2]. Graph2vec works like doc2vec by assuming that a graph is a document and the rooted subgraphs around every node in the graph are words that compose the document. Like two documents in doc2vec have similar embeddings if they consist of similar words, two graphs in graph2vec have similar embeddings if they consist of similar subgraphs, meaning that embeddings are generated in a way, both unsupervised and domain-agnostic, in which similar graphs would have similar embeddings. In the current study, signaling networks were considered undirected, so that they can be fed to the graph2vec model, and node labels are assigned as concatenated strings of the node name and the sign of the activity of each node so that the important feature of activity in a signaling network can be considered. We reason that the transformation of the graph from directed to undirected would not undermine the quality of the resulting embeddings completely, as in the case of signaling networks every connection encountered is unique for all graphs, meaning that every unique pair of nodes that exists in the dataset can have only one unique direction and sign. The graph2vec model was trained for 1 epoch and the embedding size was set to 128.

### 4 GED model

One approach to embed graphs into a high dimensional space, while maintaining the original graph-graph similarity in the high dimensional space too, is the utilization of a distance learning approach that employs Siamese encoders (shared weights) to construct graph embeddings. As proposed in the UGraphEmb framework by Bai et al., similarity or dissimilarity between graphs can be defined by domain-agnostic and unbiased distance metrics, such as Graph Edit Distance (GED), which can be used to train the model in an supervised manner [3]. One definition of GED is that of the number of operations, such as node or edge insertions and deletions, needed in order to transform one graph G1 into another graph G2 [4]. To this end, a distance learning model consisting of siamese graph convolutional encoders is trained to minimize the Mean Squared Error (MSE) between the predicted cosine distance of paired graph embeddings and the GED of the pair of input graphs. The input representation and the architecture of the encoder is similar to the one used in the deepSIBA framework [5]. The encoder consists of three graph convolutional layers, as proposed by Duvenaud et al., followed by one convolutional layer, one pooling layer and one fully connected layer, while the final graph embeddings are L2-normalized. The graphs are represented by a node matrix, containing information about the nodes' features, and edge attribute matrix, containing information about the edges' features and a connectivity matrix.

### 5 DeepSNEM-GT-MI

The deepSNEM-GT-MI model encodes the input matrices of each signaling network using two multi-head attention layers.

$$X_{prot} = \begin{bmatrix} x_{11} & x_{12} & x_{13} & \cdots & \cdots & x_{1\delta} \\ x_{21} & x_{22} & x_{23} & \cdots & \cdots & x_{2\delta} \\ \vdots & \cdots & \cdots & \cdots & \cdots & \vdots \\ x_{N1} & x_{N2} & \cdots & \cdots & \cdots & x_{N\delta} \end{bmatrix}, \text{ Nx}\delta \text{ matrix}$$

$$X_{act} = \begin{bmatrix} 0 & 1 \\ 0 & 1 \\ \vdots & \vdots \\ 1 & 0 \end{bmatrix}, \text{ Nx2 matrix}$$

$$X_{edge} = \begin{bmatrix} [0\ 1] & [1\ 0] & \cdots & [0\ 0] \\ [0\ 0] & \vdots & \vdots & \vdots \\ \vdots & \vdots & \vdots & \vdots \\ [1\ 0] & [1\ 0] & \cdots & [0\ 1] \end{bmatrix}, \text{ NxNx2.}$$

Where N is the number of nodes in a graph G and δ is the embedding size of the proteins as calculated by SeqVeq protein sequence model [6]. The edge matrix ($X_{edge}$) describes the connectivity of the graph and whether a node inhibits another node ( [0 1] ) or activates it ( [1 0] ), while because the graph is directed the inverse interaction is not defined ( [0 0] ). Additionally, before passing the inputs to the multi-head attention layer the activity matrix ($X_{act}$) is first projected linearly ($X'_{act}$) to have dimensions Nxδ as the protein matrix ($X_{prot}$) and their sum is the input feature matrix of the first multi-head attention layer.

$$X_{in} = X_{prot} + X_{act} * W_{act}$$ , where $W_{act}$ is a 2xδ matrix with trainable weights

Each multi-head attention layer computes the attention score using the key (K), query (Q) and value (V) matrices, combined with the edge matrix ($X_{edge}$) passed through a convolutional layer (new matrix E).

$$W_{attn} = softmax\left( \frac{Q * K^T}{\sqrt{\delta}} + \beta E \right)$$

Where:

- $Q = X_{in} * W_Q$
- $K = X_{in} * W_K$
- $V = X_{in} * W_V$
- $E = X_{edge} \circledast W_E + bias$

Where $W_Q$, $W_K$, $W_V$, $W_E$ and bias contain the trainable parameters.

Finally, the encoded node features of the graph (Z) are calculated by using the attention scores and the value matrix (V), which contains in a sense the encoded values of the graph, and ultimately passing through a fully connected layer :

$$\boldsymbol{X} = \boldsymbol{W_{attn}} * \boldsymbol{V}$$

$$\boldsymbol{Z} = \boldsymbol{X} * \boldsymbol{W_{FC}}$$

The output of these layers is used to produce the whole-graph representations by utilizing the Set2Set model [7]. Set2Set is a bi-directional LSTM pooling model which is utilized to summarize node embeddings (Z) into graph embeddings (s). The final node embedding size is set to 128, while the whole-graph representation embedding size is set to 256.Finally, the model is trained to maximize the mutual information between similar graphs, to ultimately create to create similar representations for similar signaling networks, while capturing the node, connectivity and edge features. The mutual information is approximated using simple discriminators (simple Artificial Neural Networks) in order to train the model using the loss described by Sun et al. in InfoGraph [8]:

$$\boldsymbol{\mathcal{L}} = -\frac{\boldsymbol{1}}{\boldsymbol{N}}\sum_{\boldsymbol{1}}^{\boldsymbol{N}} \boldsymbol{I_{\psi,\varphi}}\left(\boldsymbol{z^i}, \boldsymbol{s}\right) + \boldsymbol{\gamma D_{\varphi,\psi}}(\mathbf{Z}||\boldsymbol{U})$$

Where, $I_{\psi,\varphi}$ is the discriminator network which estimates the mutual information, while the second term corresponds to a discriminator which forces the distribution of the features of each learned graph embedding to be close to the uniform distribution.

### 6 Clustering with k-means

The deepSNEM-GT-MI embeddings were clustered using the k-means algorithm. The optimal number of clusters were selected using the elbow method. The elbow plot of the clustering is presented in Figure S1. Figure S1 shows the total within sum of squared distances between the centroids and the points of each cluster, for different values of k. We can see that the elbow starts to form around k=200. This comes in agreement with the internal diversity of the dataset, where we have 261 unique MoA labels assigned to 912 compounds. Based on the results of Figure S1, the number of clusters was set to 200.

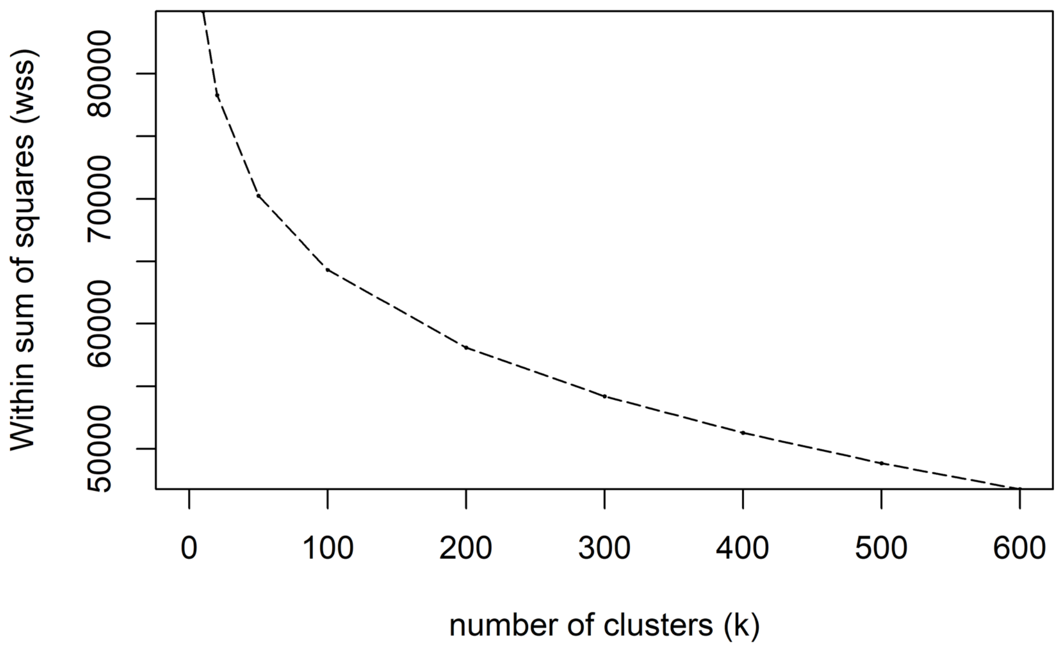


Figure S1. Elbow plot of the k-means clustering of the deepSNEM-GT-MI embeddings.

### 7 Subgraph importance

The important subgraphs for all analyzed clusters that were significantly enriched for a specific MoA are presented in Figure S2.

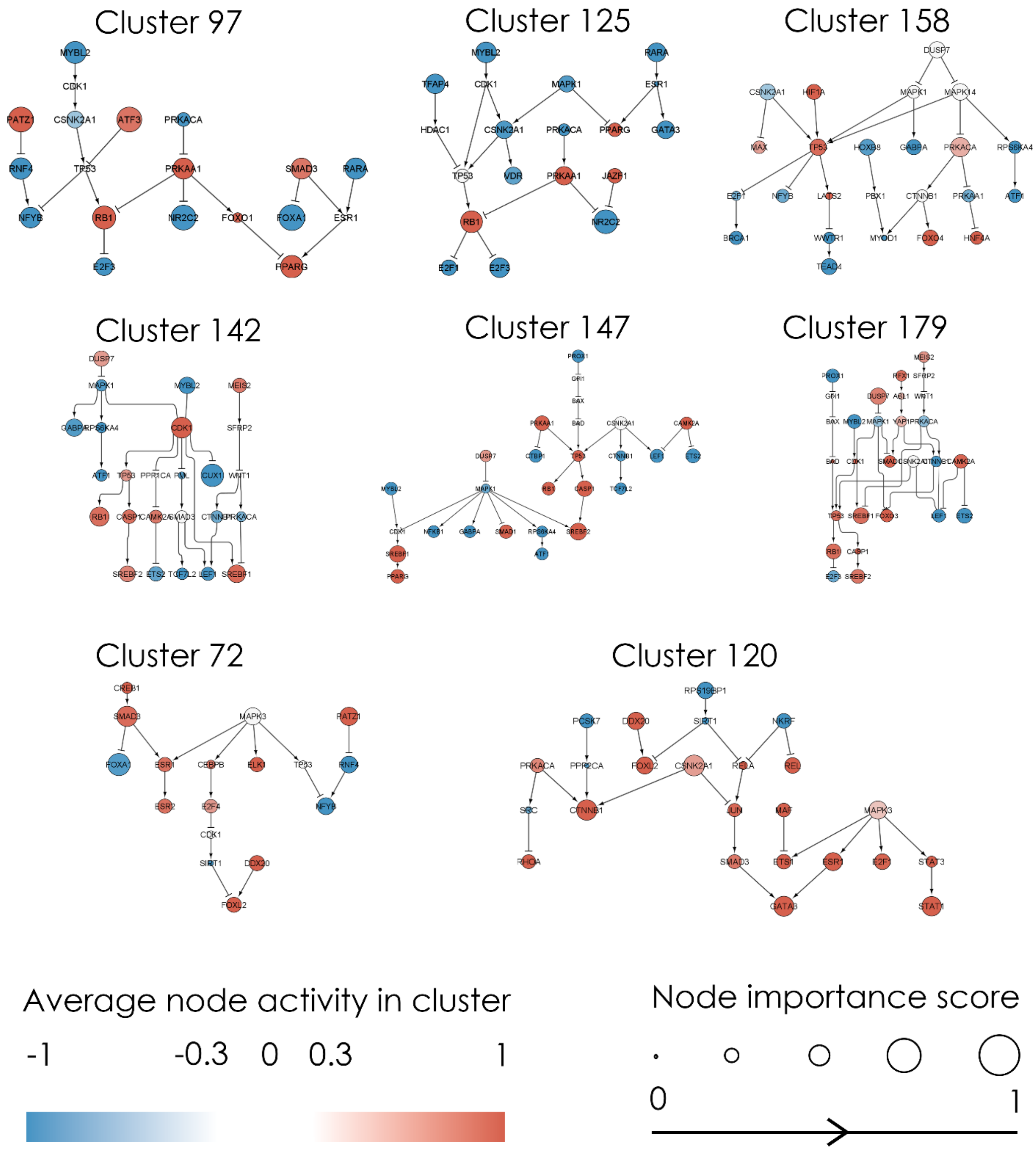


Figure S2. Important subgraphs identified for the clusters enriched for topoisomerase (Clusters 97, 125 and 158), HDAC (Clusters 142, 147 and 179), protein synthesis (Cluster 72) and ATP synthesis (Cluster 120) inhibitors. The average activity of each node in the cluster is color coded from blue to red. Blue nodes are inhibited, while red are activated. Each node's importance score, ranging from 0 to 1, is represented by the size of the node's circle.

### 8 Use case and gene-level clustering

The MicroArray gene expression profiles following compound treatment were preprocessed with the RMA algorithm, while the RNAseq data with the edgeR algorithm. The transcriptomic signatures of the CMap dataset were clustered with the k-means algorithm, similar to the signaling network embeddings. The elbow plot of the gene expression clustering is shown in Figure S3. Similar to the clustering of the deepSNEM embeddings, the number of clusters k was set to 200. Furthermore, Figure S4 shows the t-SNE projections of the gene expression profiles, where the most prevalent MoA labels in the datasets are coded with different colors.

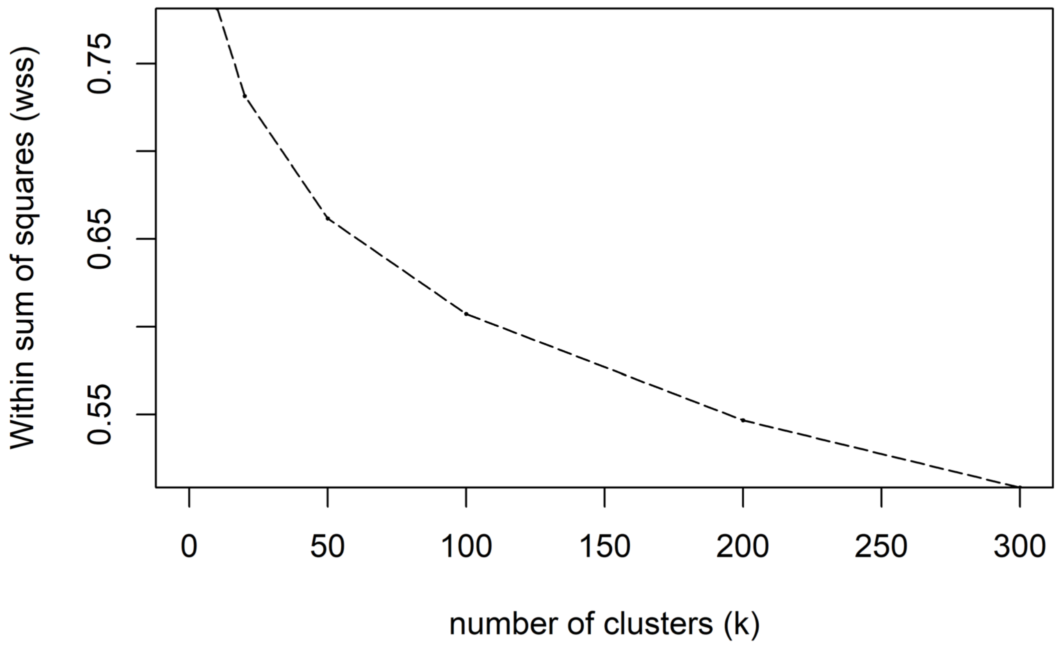


Figure S3. Elbow plot of the k-means clustering of the differential gene expression profiles.

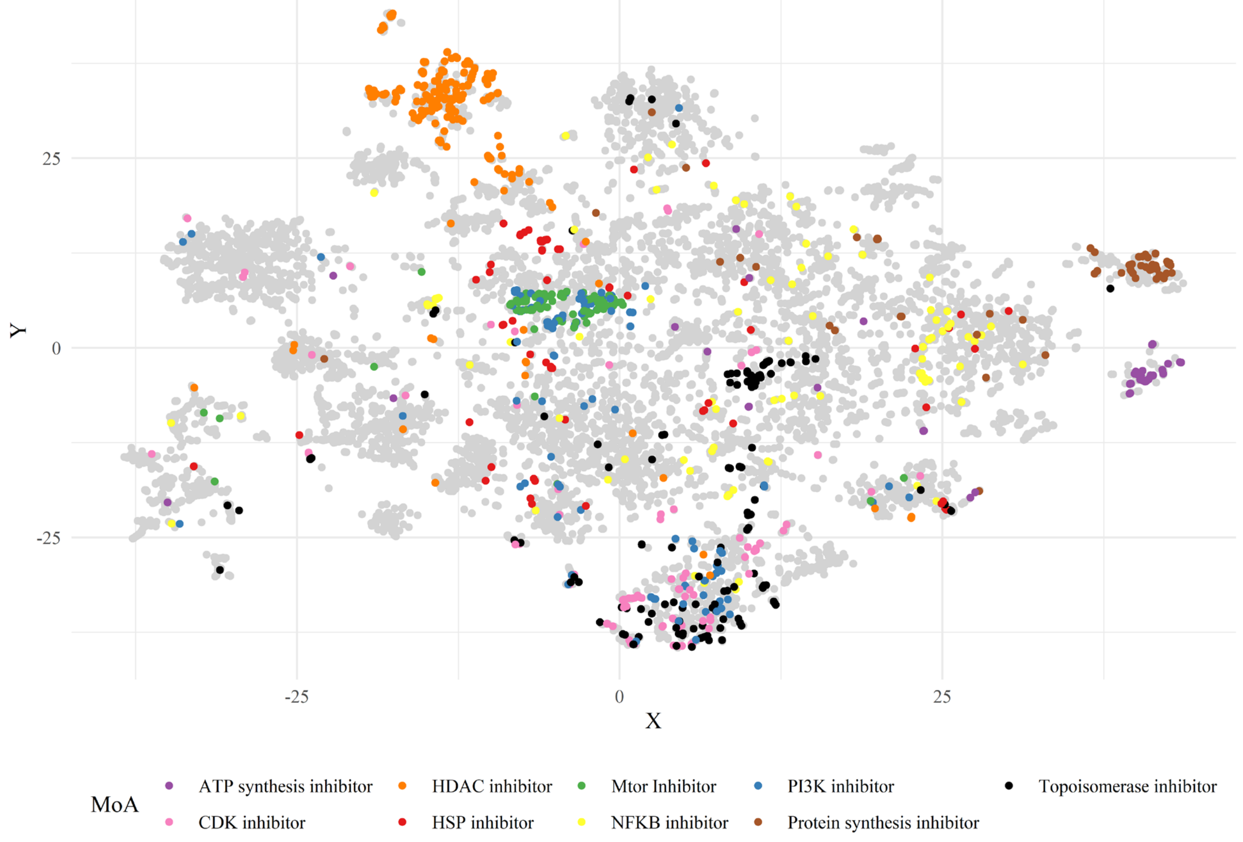


Figure S4. T-SNE projection of the gene expression profiles. Different colors represent the 9 most prevalent MoA in the dataset, while the grey color represents perturbations with either unknown or other MoA
